# Impact of Měnglà virus proteins on human and bat innate immune pathways

**DOI:** 10.1101/687400

**Authors:** Caroline G. Williams, Joyce Sweeney Gibbons, Timothy R. Keiffer, Priya Luthra, Megan R. Edwards, Christopher F. Basler

**Author notes:** equal contribution (Caroline G. Williams, Joyce Sweeney Gibbons). Corresponding Author: Center for Microbial Pathogenesis, Institute for Biomedical Sciences, Georgia State University, Atlanta, GA 30303.

## Abstract

Měnglà virus (MLAV), identified in *Rousettus* bats, is a phylogenetically distinct member of the family *Filoviridae*. Because filoviruses Ebola virus (EBOV) and Marburg virus (MARV) modulate host innate immune pathways, MLAV VP35, VP40 and VP24 proteins were compared with their EBOV and MARV homologs for innate immune pathway modulation. In human and *Rousettus* cells, MLAV VP35 behaved like EBOV and MARV VP35s, inhibiting virus-induced activation of the interferon (IFN)-β promoter. MLAV VP35 inhibited IRF3 phosphorylation and interacted with PACT, a host protein engaged by EBOV VP35 to inhibit RIG-I signaling. MLAV VP35 also inhibited PKR activation. MLAV VP40 was demonstrated to inhibit type I IFN induced gene expression in human and bat cells. It blocked STAT1 tyrosine phosphorylation induced either by type I IFN or over-expressed Jak1, paralleling MARV VP40. MLAV VP40 also inhibited virus-induced IFNβ promoter activation, a property shared by MARV VP40 and EBOV VP24. The inhibition of IFN induction was preserved in the presence of a Jak kinase inhibitor, demonstrating that inhibition of Jak-STAT signaling is not sufficient to explain inhibition of IFNβ promoter activation. MLAV VP24 did not inhibit IFN-induced gene expression or bind karyopherin α5, properties of EBOV VP24. MLAV VP24 also differed from MARV VP24 in that it failed to interact with Keap1 or activate an antioxidant response element reporter gene, due to the absence of a Keap1-binding motif. These studies demonstrate similarities between MLAV and MARV in how they suppress IFN responses and differences in how MLAV VP24 interacts with host pathways.

**Importance:** EBOV and MARV, members of the family *Filoviridae*, are highly pathogenic zoonotic viruses that cause severe disease in humans. Both viruses use several mechanisms to modulate the host innate immune response, and these likely contribute to severity of disease. Here, we demonstrate that MLAV, a filovirus newly discovered in a bat, suppresses antiviral type I interferon responses in both human and bat cells. Inhibitory activities are possessed by MLAV VP35 and VP40, which parallels how MARV blocks IFN responses. However, whereas MARV activates cellular antioxidant responses through an interaction between its VP24 protein and host protein Keap1, MLAV VP24 lacks a Keap1 binding motif and fails to activate this cytoprotective response. These data indicate that MLAV possesses immune suppressing functions that could facilitate human infection. They also demonstrate key differences in MLAV versus either EBOV or MARV engagement of host signaling pathways.

## Introduction

Měnglà virus (MLAV) was discovered when its genomic RNA was identified in the liver of a bat of the *Rousettus* genus that had been collected in Měnglà County, Yunnan Province, China (1). MLAV has been proposed to represent a new genus, *Dianlovirus*, within the family *Filoviridae*. The filovirus family includes three additional genera, *Ebolavirus*, *Marburgvirus* and *Cuevavirus*, that contain viral species isolated from or identified in mammals (2). Placement of MLAV in a distinct genus was based on its comparatively low sequence identity to other filoviruses, phylogenetic and pairwise sequence comparison (PASC) analyses (1). It was also noted to have, compared to other filoviruses, unique gene overlaps and a unique transcription start signal (1). MLAV displays some features more reminiscent of *Marburgvirus* members than *Ebolavirus* members. Specifically, MLAV RNA was identified in tissue from a *Rousettus* bat, the same genus of bat which serves as a MARV reservoir in Africa (3). In addition, the MLAV Large (L) protein exhibits closer phylogenetic relatedness to *Marburgvirus* L than to the L of other filoviruses, and like *Marburgvirus* genus members, MLAV lacks signals that direct RNA editing of the *Ebolavirus* and *Cuevavirus* glycoprotein (GP) mRNA (4).

Filoviruses are noteworthy because of their capacity to cause severe human disease (4). Some members of the *Ebolavirus* and *Marburgvirus* genera are zoonotic pathogens that have caused repeated outbreaks with substantial lethality in humans (5). The largest such outbreak on record was caused by *Zaire ebolavirus* (EBOV) and occurred in West Africa between 2013 and 2016. This resulted in upwards of 28,000 infections, more than 11,000 deaths, and the export of infected cases to the United States and Europe (6). EBOV is also the cause of the second largest filovirus outbreak, which was first recognized in August 2018 and has continued well into 2019 (7). The largest outbreak of MARV occurred in Angola between 2004-2005 and had a reported case fatality rate of 88 percent (5).

Likely contributing to the virulence of filoviruses are viral encoded proteins that target host cell innate immune signaling pathways (4). Filovirus VP35 proteins suppress interferon (IFN) α/β responses that play critical roles in innate antiviral immunity (8). VP35 impairment of IFN-α/β production occurs by inhibition of RIG-I-like receptor (RLR) signaling through several mechanisms, including VP35 binding to RLR activating dsRNAs and the interaction of VP35 with PACT, a host protein that facilitates RIG-I activation (9–20). VP35s also inhibit the phosphorylation and activation of the IFN-induced kinase PKR (21–24). EBOV VP24, but not MARV VP24, interacts with the NPI-1 subfamily of karyopherin alpha (KPNA) (also known as importin alpha) nuclear transport proteins, which includes KPNA1, KPNA5 and KPNA6 (25, 26). The NPI-1 subfamily also mediates nuclear import of STAT1 following its activation by IFN (26–28). The interaction of EBOV VP24 with KPNA competes with tyrosine phosphorylated STAT1 (pY-STAT1), blocking pY-STAT1 nuclear import and suppressing expression of IFN stimulated genes (ISGs), a response that mediates the antiviral effects of IFN (25, 26, 29, 30). MARV VP40 protein has been demonstrated to suppress IFN-induced signaling and ISG expression, while EBOV VP40 has no known role in IFN antagonism (31). Activation of the Jak family of kinases associated with IFN receptors is inhibited by MARV VP40, blocking phosphorylation and activation of the downstream STAT proteins, including STAT1 (31–33). EBOV VP24 and MARV VP40 have also been described to modestly inhibit IFN-α/β production, although the mechanism(s) are not defined (34, 35). While MARV VP24 does not appear to block IFN responses, it has been demonstrated to interact with Kelch-like ECH-associated protein 1 (Keap1). Under homeostatic conditions, Keap1, a cellular substrate adaptor protein of the Cullin3/Rbx1 ubiquitin E3 ligase complex, targets the transcription factor Nuclear factor erythroid 2-related factor 2 (Nrf2) for polyubiquitination and proteasomal degradation (36–38). MARV VP24 disrupts the Keap1-Nrf2 interaction, leading to Nrf2-induced expression of genes possessing antioxidant response elements (ARE) (36–38). This activity induces a cytoprotective state that may prolong the life of MARV infected cells. MARV VP24 also relieves Keap1 repression of the NF-κB pathway (39).

Given the link between EBOV and MARV innate immune suppressors and virulence, and the unknown potential of MLAV to cause human disease, this study sought to determine whether MLAV possesses effective suppressors of innate immunity. Given the differences in innate immune evasion mechanisms between EBOV and MARV, it was also of interest to determine whether MLAV innate immune evasion mechanisms more closely resemble EBOV or MARV. The data demonstrate that MLAV VP35 functions as an IFN antagonist by mechanisms that mirror those of EBOV and MARV VP35. MLAV VP40 is demonstrated to act as a suppressor of IFN-induced signaling, whereas MLAV VP24 does not, mirroring the inhibitory functions of MARV. Both MLAV VP35 and VP40 effectively suppressed IFN responses in human and *Rousettus* cells. Interestingly, MLAV VP24 does not detectably interact with Keap1 or activate ARE gene expression due to the absence of Keap1-binding sequences found in MARV VP24. Cumulatively, the data demonstrate the presence of IFN evasion functions in MLAV that are effective in human cells, suggesting the virus may have the capacity to cause human disease. The similarities in VP40 immune evasion functions are consistent with a closer genetic relationship of MLAV to MARV than EBOV, but the differences in VP24 function are consistent with MLAV occupying a distinct genus within the filovirus family.

## Material and Methods

### Cells and viruses

HEK293T cells were maintained in Dulbecco’s Modified Eagle Medium (DMEM), supplemented with 10% fetal bovine serum (FBS) and cultured at 37°C and 5% CO_2_. RO6E cells, immortalized fetal cells from *Rousettus aegyptiacus*, were obtained from BEI Resources and maintained in DMEM F12 and supplemented with 5% FBS. Sendai Virus Cantell (SeV) was grown in 10-day-old embryonating chicken eggs for forty-eight hours at 37°C.

### Plasmids

MLAV VP35, VP40, and VP24 coding sequences (based on accession number KX371887) were synthesized by Genscript. The synthesized open reading frames were cloned into a pCAGGS expression vector with a FLAG-tag at the N-terminus of each coding sequence. EBOV and MARV viral proteins, GFP-STAT1, HA-Jak1, HA-PACT, HA-KPNA5, HA-Keap1 and IRF3 expression plasmids were previously described(8, 13, 25, 31, 36). VP24 K-loop chimeras were made using overlapping PCR. MARV VP24 residues 202-RRIDIEPCCGETVLSESV-219 were inserted into MLAV VP24 between residues 202 and 219 (MLAV VP24_MARV 202-219_) and the corresponding MLAV residues 202-RAINASGRENESVVQNPI-219 were inserted into MARV VP24 at the same position (MARV VP24_MLAV 202-219_).

### Cytokines

Universal type I IFN (UIFN) (PBL) was used at 1000 U/mL in DMEM supplemented with 0.3% FBS for 30 minutes at 37°C, unless otherwise stated.

### Phosphorylation assays

HEK293T cells were transfected using Lipofectamine 2000^®^ (Life Technologies). Twenty-four hours post transfection, cells were mock-treated, UIFN-treated or SeV-infected, depending on the assay. Subsequently, cells were lysed in NP40 buffer (50mM Tris-HCl [pH 8.0], 280mM NaCl, 0.5% NP-40) supplemented with cOmplete™ protease inhibitor cocktail (Roche) and PhosSTOP (Roche). Lysates were incubated for ten minutes on ice and clarified for ten minutes at 21,100 x g at 4°C. Phosphorylation status of the proteins was determined by western blot.

### IFNβ− and ISG54-promoter reporter gene assays

HEK293T cells (5×10^4^) and RO6E cells (2×10^5^) were co-transfected with 25 ng of an IFNβ promoter-firefly luciferase reporter plasmid or an interferon stimulated gene 54 (ISG54) promoter-firefly luciferase reporter plasmid, 25 ng of a constitutively expressing *Renilla* luciferase plasmid (pRLTK, Promega) and the indicated viral protein expression plasmids – HEK293T cells: 62.5, 6.25, and 0.625 ng for VP35 and VP40 and 25, 2.5, and 0.25 ng for VP24; RO6E cells: 250, 25, and 2.5 ng for EBOV and MARV proteins and 125, 12.5, and 1.25 ng for MLAV proteins. Twenty-four hours post transfection cells were mock-treated, SeV-infected (150 hemagglutinin activity units (HAU)) or UIFN-treated (1000 U/mL). Eighteen hours post-infection or treatment, cells were lysed and analyzed for luciferase activity using a Dual Luciferase^®^ Reporter Assay System (Promega) per the manufacturer’s protocol. Firefly luciferase activity was normalized to *Renilla* luciferase activity. Assays were performed in triplicate; error bars indicate the standard error of the mean (SEM) for the triplicate. Viral protein expression was confirmed by western blot.

### IFNβ reporter gene assay in the presence of a Jak1/Jak2 inhibitor

HEK293T cells (5×10^4^) were co-transfected with 25 ng of an IFNβ promoter-firefly luciferase reporter plasmid, 25 ng of pRLTK *Renilla* luciferase reporter plasmid and 62.5, 6.25, and 0.625 ng of the indicated viral protein expression plasmids. Twenty-four hours post-transfection, cells were pre-treated for one hour with 5μM of Ruxolitinib (SelleckChem), a Jak1/Jak2 inhibitor, and then mock- or SeV-infected in the presence of the inhibitor (40). Eighteen hours post-infection or treatment, cells were lysed and assayed using a dual luciferase assay and analyzed as above. To verify inhibition of Jak1/Jak2 by Ruxolitnib, cells were transfected with 25 ng of an ISG54 promoter-firefly luciferase reporter plasmid and 25 ng of pRLTK reporter plasmid. Twenty-four hours post-transfection, cells were pre-treated for one hour with 5μM of Ruxolitinib, and then mock- or UIFN-treated for eighteen hours in the presence of the inhibitor and assayed for luciferase activity as above.

### ARE reporter assay

HEK293T cells (5×10^4^) were co-transfected with an antioxidant response element (ARE) reporter gene, pGL4.37 [luc2P/ARE/Hygro] (Promega) (30 ng) and a pRLTK reporter plasmid (25 ng) along with either empty vector or 62.5, 6.25, and 0.625 ng of EBOV, MARV, MLAV VP24 or chimeric MARV and MLAV expression plasmids. Eighteen hours post-transfection, luciferase activity was assessed and analyzed as above.

### Co-immunoprecipitation assays

HEK293T cells were co-transfected with plasmids for FLAG-tagged MLAV proteins, HA-tagged host proteins, and pCAGGS empty vector using Lipofectamine 2000^®^ (LifeTechnologies). Twenty-four hours post-transfection cells were rinsed with PBS and lysed in NP40 buffer supplemented with cOmplete™ protease inhibitor cocktail (Roche). Lysates were clarified by centrifugation and incubated with anti-FLAG M2 magnetic beads (Sigma-Aldrich) for two hours at 4°C. Beads were washed 5 times in NP40 buffer and precipitated proteins were eluted by boiling with SDS sample loading buffer or elution with 3X FLAG peptide (Sigma-Aldrich). Whole cell lysates and immunoprecipitated samples were analyzed by western blot.

### Western blot analysis

Blots were probed with anti-FLAG (Sigma-Aldrich), anti-β-tubulin (Sigma-Aldrich), anti-HA (Sigma-Aldrich), anti-phospho-IRF3 (S396) (Cell Signaling), anti-IRF3 (Santa Cruz), anti-phospho-STAT1 (Y701) (BD Transduction Laboratories), anti-STAT1 (BD Transduction Laboratories), anti-phospho-PKR (T446) (ABCAM), or anti-PKR (Cell Signaling) antibodies, as indicated. Antibodies were diluted in Tris-buffered saline with 0.1% Tween-20 (TBS-T) with 5% milk or, to detect phospho-proteins, 5% bovine serum albumin.

### VP40 Budding Assay

10 µg of MARV and MLAV VP40 expression plasmids were transfected into either HEK293T (3.0×10^6^) or RO6E (1.2×10^6^) cells using Lipofectamine 2000^®^ (LifeTechnologies). Media was harvested 48 hours post-transfection, briefly clarified by centrifugation, and layered over a 20% sucrose cushion in NTE buffer (10 mM NaCl, 10 mM Tris [pH 7.5], 1 mM EDTA [pH 8.0]). The samples were then subjected to ultracentrifugation in a Beckman SW41 rotor at 222,200 x g for 2 hours at 10°C; media was aspirated after ultracentrifugation and virus-like particles (VLPs) were solubilized in NTE buffer at 4°C overnight. Cellular lysates were generated by washing transfected cells with PBS and lysing cells in NP40 buffer containing cOmplete™ protease inhibitor cocktail (Roche). To detect presence of VP40, 5% of cell lysates and 10% of VLPs were visualized by western blotting.

### Statistics

Statistical significance was determined by one-way ANOVA followed with Tukey multiple comparison as compared to the indicated control; **p < 0.0001, * p < 0.001 (GraphPad PRISM8).

## Results

### MLAV VP35 blocks virus induced IFNβ promoter activation in both human and bat cells

As a measure of the capacity of MLAV VP35, VP40 and VP24 to modulate type I IFN production, the human cell line HEK293T or the *Rousettus* bat cell line RO6E were assessed by reporter gene assay for their effect on Sendai virus (SeV) induced IFNβ promoter activation. Either empty vector or FLAG-tagged expression plasmids for the VP35, VP40 and VP24 proteins of EBOV, MARV and MLAV were co-transfected with an IFNβ promoter firefly luciferase reporter and a constitutively expressing *Renilla* luciferase plasmid. Twenty-four hours post-transfection, cells were either mock-infected or infected with SeV, a potent activator of the IFNβ promoter (41, 42). As expected, SeV infection activated the IFNβ promoter in the absence of viral protein expression. EBOV and MARV VP35 impaired IFNβ reporter activation in a dose-dependent manner in both cell lines, with EBOV exhibiting greater potency (Figure 1A and 1B). Similarly, MLAV VP35 dramatically diminished IFNβ promoter activity in a dose dependent manner (Figure 1A and 1B).

**Figure 1.**
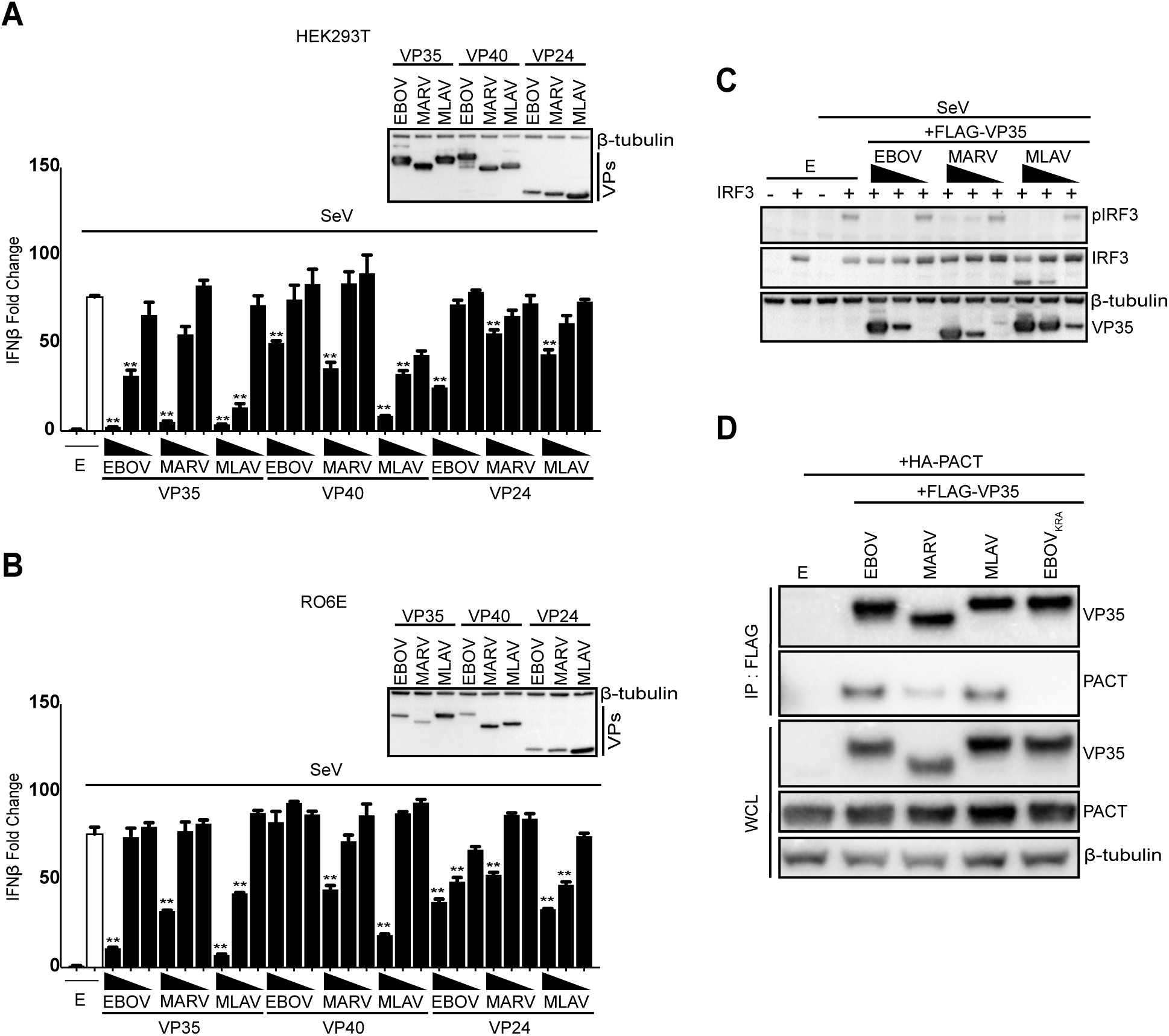
MLAV VP35 blocks virus-induced IFNβ promoter activation in both human and bat cells. **(A)** HEK293T cells were transfected with an IFNβ promoter-firefly luciferase reporter plasmid, a constitutively-expressed *Renilla* luciferase reporter plasmid and either empty vector (E) or the specified FLAG-tagged viral proteins. The concentration of VP35 and VP40 plasmids were 62.5 ng, 6.25 ng and 0.625 ng; the concentration of VP24 plasmids were 25 ng, 2.5 ng and 0.25 ng. Twenty-four hours post-transfection, cells were either mock or Sendai virus (SeV)-infected. Firefly and *Renilla* luciferase activities were determined eighteen hours post-infection using a dual luciferase assay. Fold induction was determined relative to the vector only, mock-infected samples. Assays were performed in triplicate, error bars represent the SEM for the triplicate. Whole cell lysates (WCL) were analyzed by western blot with anti-FLAG and anti-β-tubulin antibodies. **(B)** RO6E cells were assayed as described above, except the concentration of EBOV and MARV VP35, VP40 and VP24 plasmids were 250 ng, 25 ng and 2.5 ng and the concentration of MLAV VP35, VP40 and VP24 plasmids were 125 ng, 12.5 ng and 1.25 ng. **(A-B)** Statistical significance was determined by performing a one-way ANOVA followed with Tukey multiple comparison as compared to SeV-infected control (white bar); **p < 0.0001, * p < 0.001. VPs – viral proteins. **C)** HEK293T cells were transfected with empty vector (E) or IRF3 expression plasmid, as indicated, and FLAG-tagged EBOV, MARV, MLAV VP35. The concentration of VP35 plasmids were 2,000 ng, 400 ng and 80 ng. Cells were either mock or SeV-infected for four hours. Whole cell lysates (WCL) were analyzed by western blot with anti-pIRF3 (S396), anti-total IRF3, anti-FLAG (VP35), and anti-β-tubulin antibodies. **(D)** HEK293T cells were transfected with empty vector (E), or plasmids that express FLAG-tagged EBOV VP35, MARV VP35, MLAV VP35, or dsRNA binding mutant EBOV VP35_KRA_ and HA-tagged PACT, as indicated. Immunoprecipitations (IP) were performed with anti-FLAG antibody. Western blots were performed for detection of VP35 (anti-FLAG antibody), PACT (anti-HA antibody), and β-tubulin.

Expression of EBOV VP24, Lloviu virus (LLOV) VP24 or MARV VP40 has also been reported to impair IFNβ and, in the case of EBOV VP24, IFNλ production (34, 35, 43). In the present study, in HEK293T cells, modest inhibition of IFNβ promoter activation was evident for EBOV VP24, EBOV VP40, and MARV VP40. MLAV VP40 exhibited potent dose-dependent inhibition of IFNβ promoter activation (Figure 1A). Weak, but statistically significant inhibition of IFNβ reporter gene expression was detected for MARV VP24 and MLAV VP24, however, this minimal inhibition may not be biologically relevant. In RO6E cells, MLAV VP40 inhibition of IFNβ promoter activation was also detected but only at the highest concentration of transfected plasmid (Figure 1B).

EBOV and MARV VP35 inhibition of RLR signaling pathways results in inhibition of the phosphorylation and activation of transcription factor interferon regulatory factor 3 (IRF3) (9, 44, 45). In order to determine whether MLAV VP35 can inhibit activation of IRF3, HEK293T cells were co-transfected with either empty vector or an IRF3 expression plasmid and plasmids that express FLAG-tagged EBOV, MARV, and MLAV VP35 (Figure 1C). Twenty-four hours post-transfection, cells were either mock- or SeV-infected to induce IRF3 phosphorylation. Over-expression of IRF3 substantially increased detection of the phosphorylated form. As previously reported, EBOV VP35 potently inhibited IRF3 phosphorylation. MARV VP35 also inhibited IRF3 phosphorylation, although less efficiently, consistent with less robust inhibition of RIG-I signaling as compared to EBOV VP35 (8). MLAV VP35 inhibited IRF3 phosphorylation comparable to EBOV VP35.

EBOV and MARV VP35 interact with host protein PACT, and this interaction contributes to VP35 inhibition of RIG-I signaling (13, 23). To determine if MLAV VP35 suppresses IFN production through a similar mechanism, the PACT-VP35 interactions were evaluated by co-immunoprecipitation assay (Figure 1D). FLAG-tagged EBOV, MARV, and MLAV VP35 or empty vector expression plasmids were co-transfected with HA-tagged PACT in HEK293T cells. A VP35 dsRNA binding mutant (VP35_KRA_) that has previously been previously shown to lack the ability to interact with PACT was included as a negative control(13). All three wildtype VP35 proteins were demonstrated to interact with PACT, with MLAV VP35 interacting comparably to EBOV VP35. Together, these data suggest that MLAV VP35 employs mechanisms similar to EBOV and MARV VP35 for inhibition of RIG-I dependent activation of type I IFN responses and that the potency of inhibition is similar to EBOV VP35.

### MLAV VP35 protein inhibits phosphorylation of PKR in human cells

To assess whether MLAV VP35 can inhibit activation of PKR, HEK293T cells were transfected with FLAG-tagged EBOV, MARV, and MLAV VP35, or empty vector expression plasmids. Consistent with previous literature, EBOV VP35 and MARV VP35 inhibited SeV-induced PKR phosphorylation (Figure 2). MLAV VP35 also inhibited activation of PKR in a concentration dependent manner (Figure 2).

**Figure 2.**
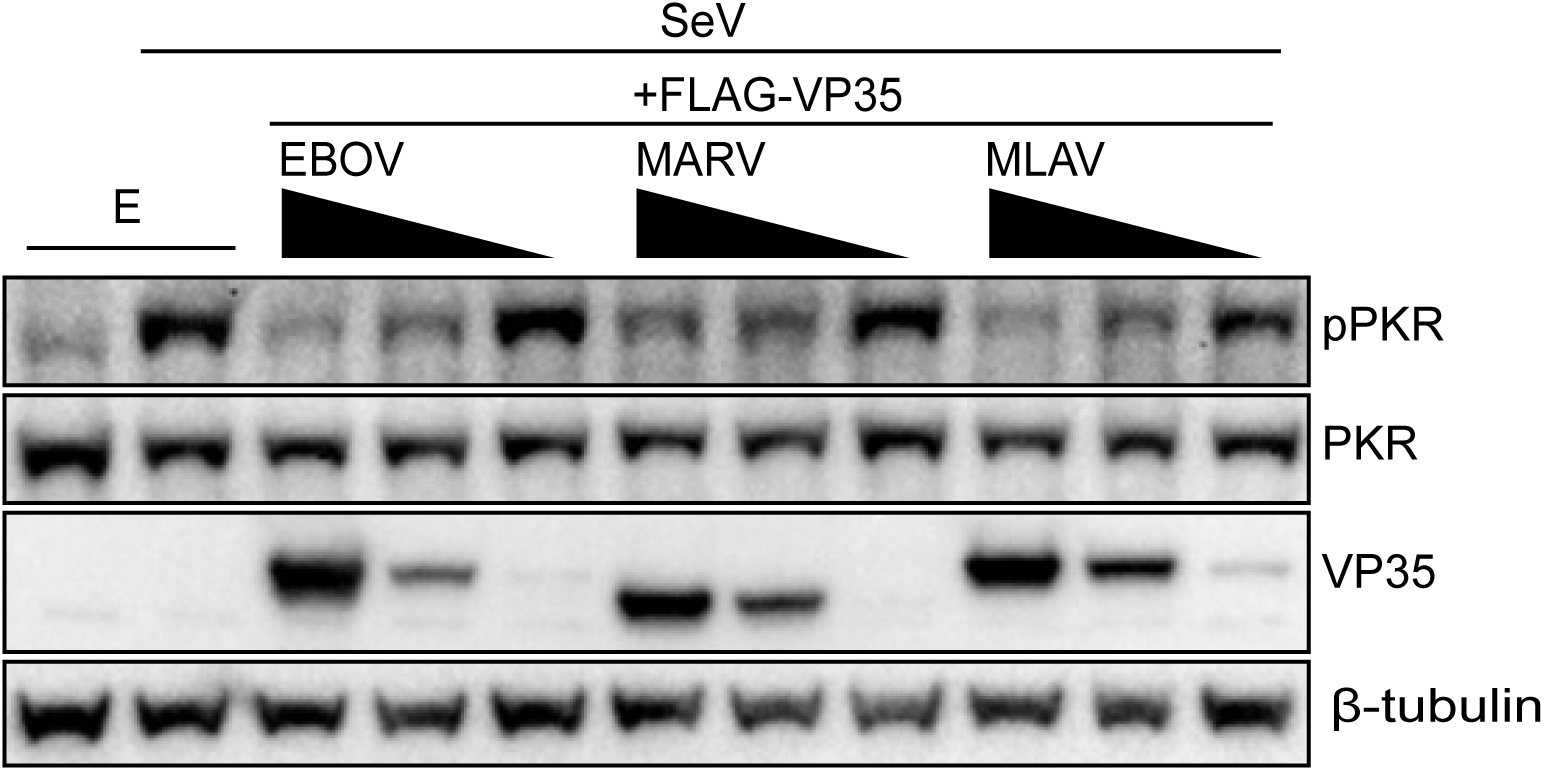
MLAV VP35 inhibits virus-induced PKR activation. HEK293T cells were transfected with empty vector (E) or expression plasmids for FLAG-tagged EBOV, MARV and MLAV VP35, as indicated. The concentration of VP35 plasmids were 2,000 ng, 400 ng and 80 ng. Twenty-four hours post-transfection, cells were mock- or SeV-infected. Eighteen hours post infection, whole cell lysates (WCL) were assessed by western blot for levels of total and phosphorylated PKR using anti-FLAG (VP35), anti-total PKR, anti-pPKR (T446) and anti-β-tubulin antibodies.

### MLAV VP40 protein inhibits responses to type I IFN in both human and bat cells

To test the effects of MLAV VP35, VP40 and VP24 on the response of cells to exogenous type I IFN, empty vector or expression plasmids for FLAG-tagged VP35, VP40 and VP24 proteins of EBOV, MARV and MLAV were co-transfected with an IFN-responsive ISG54 promoter-firefly luciferase reporter plasmid and a plasmid that constitutively expresses *Renilla* luciferase. Twenty-four hours post-transfection, cells were either mock- or type I IFN-treated. The ISG54 reporter was activated by IFN-treatment in the absence of viral protein expression, albeit less so in the bat cells (Figure 3A and 3B). As expected, both MARV VP40 and EBOV VP24 strongly inhibited ISG54 reporter activity in both human and bat cell lines (Figure 3A and 3B). Similar to MARV VP40, MLAV VP40 demonstrated potent inhibition of the ISG54 reporter in both cell types. MLAV VP24 did not substantially inhibit IFN-induced expression and minor decreases in reporter gene activity were also seen with EBOV VP40 and MARV VP24, however, the effects were minimal suggesting that this inhibition may not be biologically relevant.

**Figure 3.**
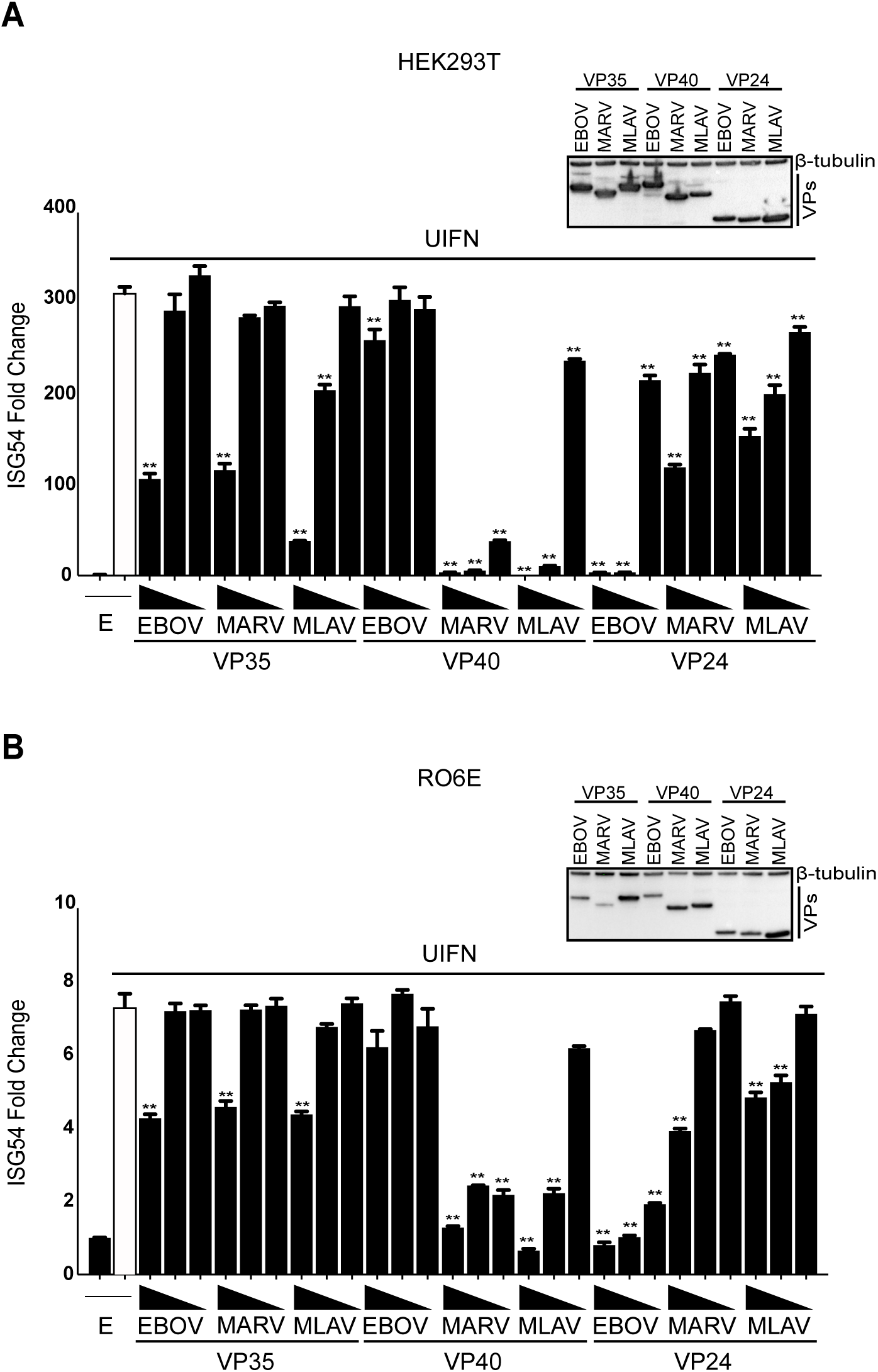
MLAV VP40 protein inhibits responses to type I IFN in both human and bat cells. **(A)** HEK293T cells were transfected with an ISG54 promoter-firefly luciferase reporter plasmid, a constitutively-expressed *Renilla* luciferase reporter plasmid and either empty vector (E) or the specified FLAG-tagged viral proteins. The concentration of VP35 and VP40 plasmids were 62.5 ng, 6.25 ng and 0.625 ng; the concentration of VP24 plasmids were 25 ng, 2.5 ng and 0.25 ng. Twenty-four hours post-transfection, cells were either mock or UIFN treated. Eighteen hours post-treatment, firefly and *Renilla* luciferase activities were determined. Firefly luciferase values were normalized to *Renilla* luciferase values and fold induction was calculated relative to the vector only, mock-treated samples. Experiments were performed in triplicate, error bars represent the SEM for the triplicate. Whole cell lysates (WCL) were analyzed by western blot with anti-FLAG and anti-β-tubulin antibodies. **B)** RO6E cells were transfected as described above, except the concentration of EBOV and MARV VP35, VP40 and VP24 plasmids were 250 ng, 25 ng and 2.5 ng and the concentration of MLAV VP35, VP40 and VP24 plasmids were 125 ng, 12.5 ng and 1.25 ng. **(A-B)** Statistical significance was determined by performing a one-way ANOVA followed with Tukey’s multiple comparison test as compared to UIFN-treated control (white bar); **p < 0.0001, *p < 0.001.

MARV VP40 has been shown to be a potent inhibitor of IFN-α/β induced phosphorylation of STAT1; whereas EBOV VP24 inhibits this pathway by blocking nuclear transport of pY-STAT1(25, 26, 30). To determine whether inhibition of IFN responses is due to inhibition of STAT1 phosphorylation, HEK293T cells were co-transfected with empty vector or expression plasmids for FLAG-tagged EBOV, MARV, and MLAV VP24 or VP40. GFP-STAT1 was (Figure 4A) or was not (Figure 4B) included in the transfection. Addition of IFN triggered the phosphorylation of GFP-STAT1 and endogenous STAT1 in the vector only samples. Among the EBOV and MARV constructs, only MARV VP40 was inhibitory. MLAV VP40 also inhibited STAT1 tyrosine phosphorylation to a similar degree as MARV VP40. MLAV VP24 did not detectably affect STAT1 phosphorylation.

**Figure 4.**
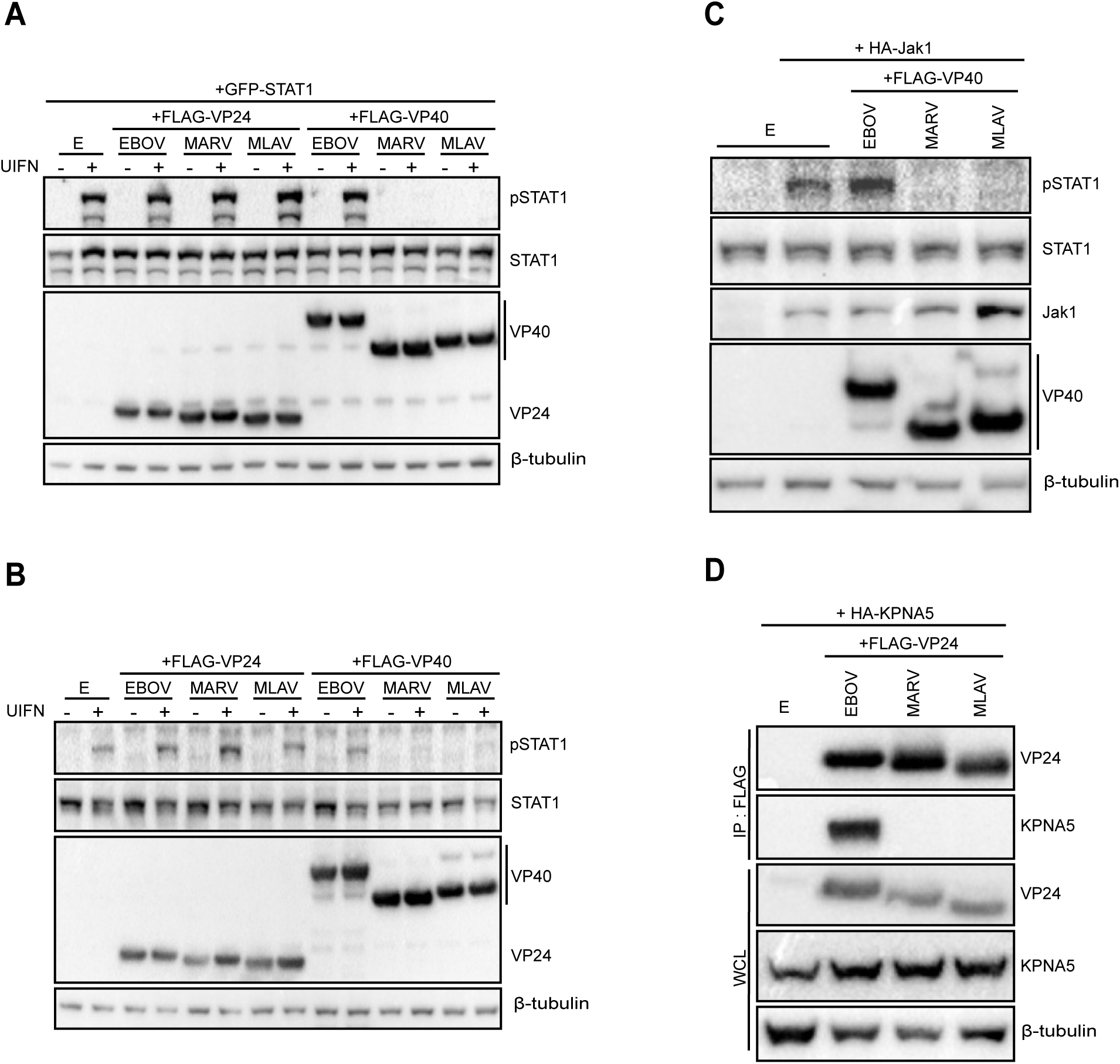
MLAV VP40 protein inhibits type I IFN induced gene expression and Jak-STAT signaling. HEK293T cells were transfected with empty vector (E), FLAG-tagged VP24s or VP40s from EBOV, MARV and MLAV, as indicated. Twenty-four hours post-transfection, cells were treated with UIFN for 30 minutes and the phosphorylation status of exogenous GFP-STAT1 **(A)** or endogenous STAT1 **(B)** was assessed by western blotting. **(C)** HEK293T cells were co-transfected with empty vector (E) or FLAG-tagged VP40s from EBOV, MARV and MLAV and HA-tagged Jak1 expression plasmids. Twenty-four hours post-transfection cells were lysed and phosphorylation status of endogenous STAT1 was analyzed by western blot with anti-FLAG, anti-STAT1, anti-pSTAT1 (Y701), and anti-β-tubulin antibodies. **(D)** HEK293T cells were co-transfected with FLAG-tagged EBOV, MARV, MLAV VP24, and HA-tagged KPNA5. Immunoprecipitation (IP) was performed with anti-FLAG antibody and precipitates and whole cell lysates (WCL) were assessed by using anti-FLAG (VP24), anti-HA (KPNA5) and anti-β-tubulin antibodies.

MARV VP40 inhibits STAT1 phosphorylation following over-expression of Jak1(31). To determine whether MLAV VP40 can prevent Jak1 induced STAT1 phosphorylation, HA-tagged Jak1 was co-transfected with empty vector or FLAG-tagged EBOV, MARV or MLAV VP40. As expected, expression of exogenous Jak1 induced STAT1 tyrosine phosphorylation, and this was suppressed in the presence of MARV VP40 (Figure 4C). Similarly, MLAV VP40 prevented Jak1-dependent STAT1 phosphorylation, suggesting that MLAV VP40 inhibits IFN signaling through the same mechanism used by MARV VP40.

EBOV VP24 interacts with NPI-1 subfamily members of the KPNA nuclear transporters, including KPNA5, to block nuclear import of pY-STAT1(26–28). To assess whether MLAV VP24 interacts with KPNA5, co-immunoprecipitation assays were performed in HEK293T cells (Figure 4D). KPNA5 did not precipitate in the absence of a co-expressed protein. Among FLAG-tagged EBOV, MARV, and MLAV VP24, only EBOV VP24 detectably interacted with KPNA5. The absence of MLAV VP24-KPNA5 interaction is consistent with the inability of MLAV VP24 to inhibit IFN-induced gene expression.

### MLAV and MARV VP40 bud with similar efficiencies from human and bat cells

Filovirus VP40 proteins play a critical role in budding of new virus particles, and expression of VP40 is sufficient for formation and budding of VLPs (46–50). Given the functional similarities of MLAV and MARV VP40s to inhibit IFN responses, it was of interest to compare budding capacity. Upon expression in human and bat cells, both MLAV and MARV VP40 expressed to similar levels and budded from human and bat cells to a similar extent. (Figure 5A and 5B).

**Figure 5.**
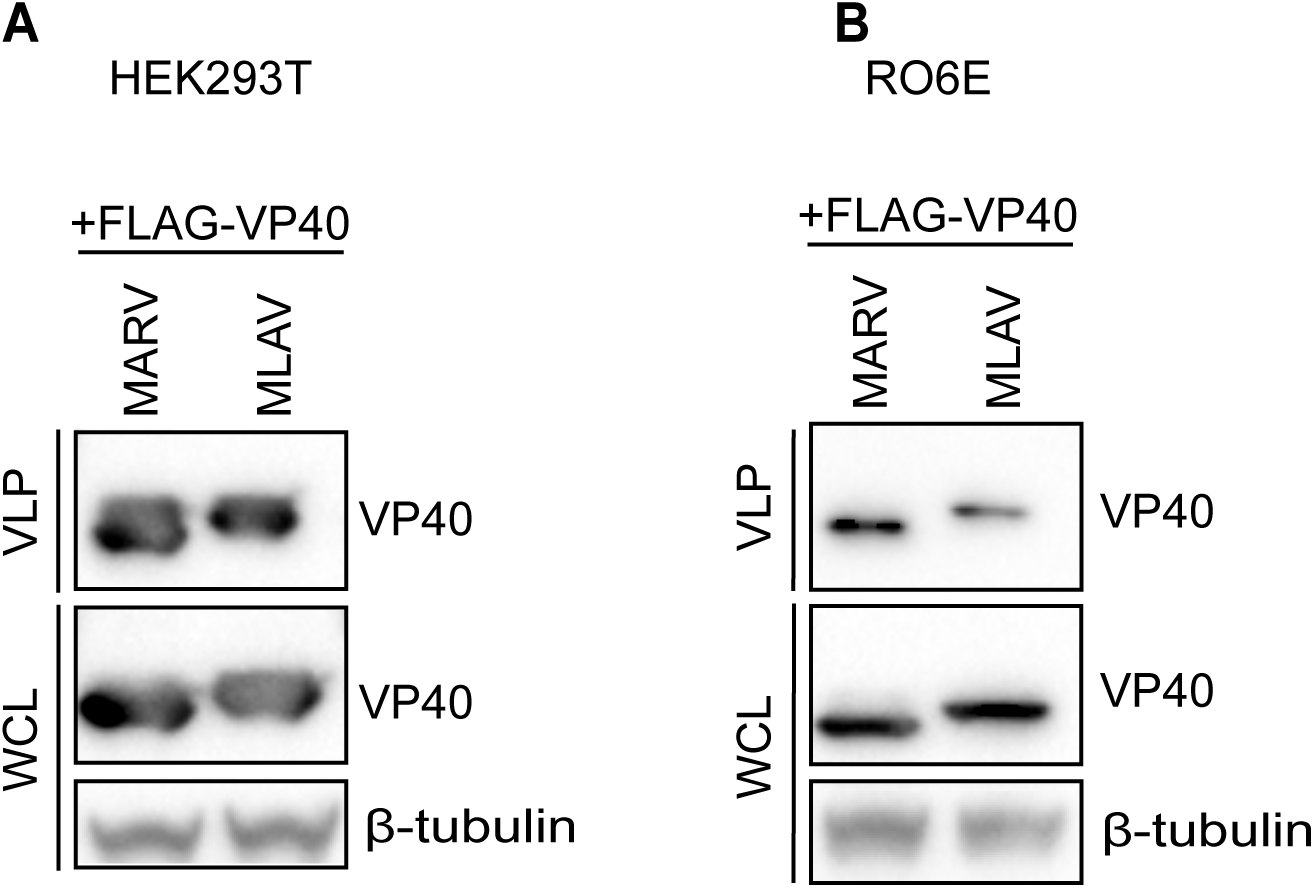
MLAV VP40 is capable of budding from both human and bat cells. To compare the budding of EBOV, MARV, and MLAV VP40 proteins from different cell lines, VLP assays were performed in a HEK293T cells **(A)** and RO6E cells **(B)**, as previously described. Presence of VP40 in VLPs and whole cell lysates (WCL) were determined by western blot using anti-FLAG antibody.

### MLAV VP40 and EBOV VP24 inhibition of IFNβ promoter activation occurs independently of Jak-STAT signaling

The type I IFN response includes a positive feedback loop whereby secreted IFN upregulates pattern recognition receptors, such as RIG-I and transcription factors such as IRF7, to amplify the response (51). It was therefore of interest to test the hypothesis that MLAV VP40, MARV VP40 and EBOV VP24 inhibit virus-induced induction of the IFN response as a result of their inhibition of IFN-induced positive feedback loop. Activation of the IFNβ promoter by SeV was therefore assessed by reporter gene assay in the absence or presence of the Jak1/Jak2 inhibitor Ruxolitinib. In this experiment, cells were transfected with empty vector or FLAG-tagged expression plasmids for the EBOV VP35, EBOV, MARV and MLAV VP40 and EBOV VP24, pre-treated with DMSO or Ruxolitinib and then mock- or SeV-infected, in the absence or presence of the inhibitor (Figure 6A). EBOV VP35 acted as a potent suppressor of IFNβ promoter activation under these conditions. EBOV VP40, MARV VP40, MLAV VP40, and EBOV VP24 all suppressed IFNβ promoter activation to similar extents in the absence or presence of the Jak kinase inhibitor. To confirm that inhibition of IFN induced signaling was complete, cells transfected with an ISG54 promoter reporter gene were DMSO or Ruxolitinib treated and then mock or IFN-treated. As expected, IFN activated the ISG54 promoter in the presence of DMSO but not Ruxolitinib (Figure 6B). These data suggest that EBOV VP40, MARV VP40, MLAV VP40 and EBOV VP24 all utilize an additional mechanism, outside of inhibition of STAT1, to impair IFN induction.

**Figure 6.**
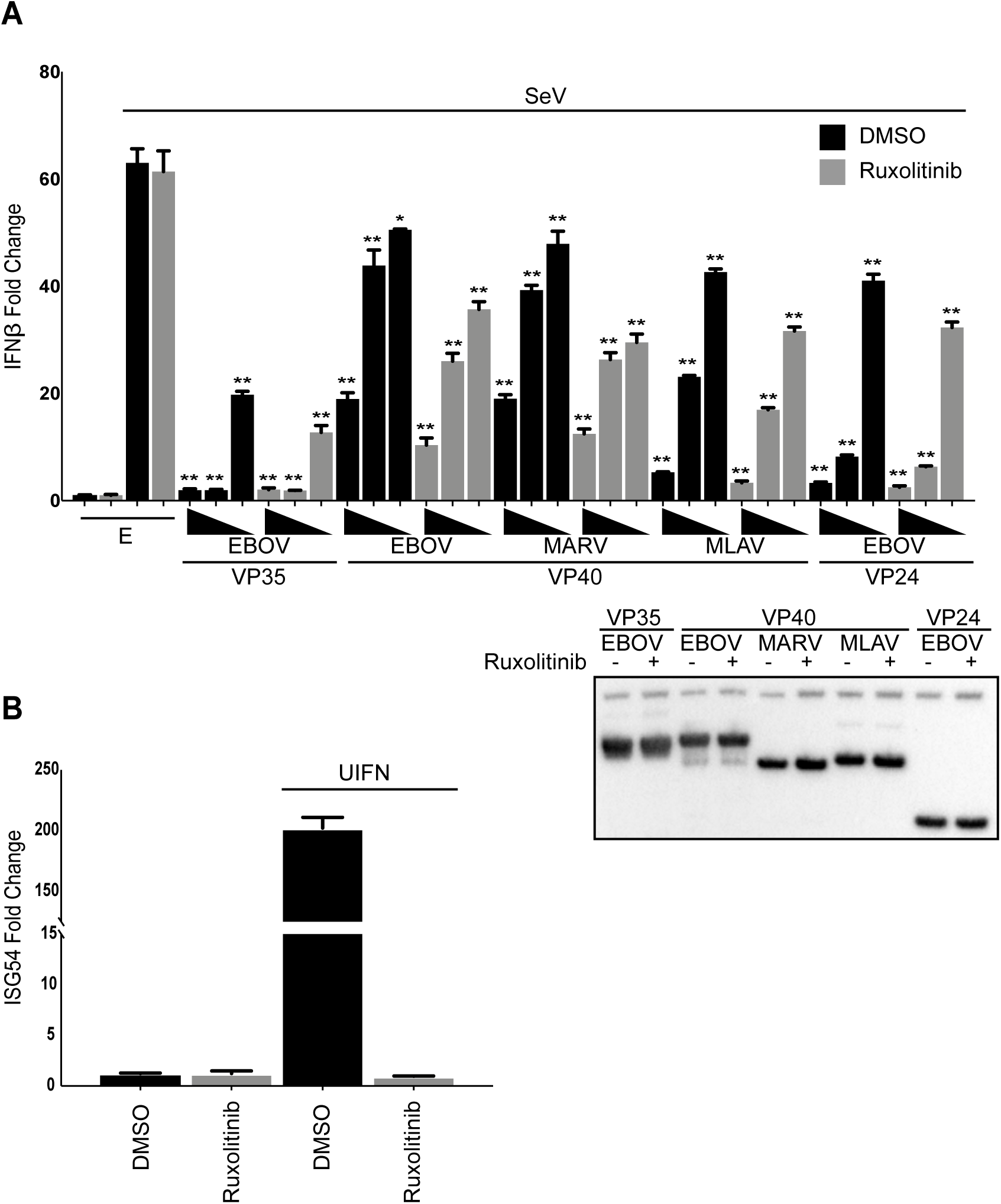
MLAV VP40 blocks virus-induced IFNβ promoter activation independently of Jak-STAT signaling. **(A)** HEK293T cells were transfected with an IFNβ promoter-firefly luciferase reporter plasmid, a constitutively-expressed *Renilla* luciferase reporter plasmid and either empty vector (E) or the specified FLAG-tagged viral proteins. The concentration of VP35, VP40, and VP24 plasmids were 62.5 ng, 6.25 ng, and 0.625 ng. Twenty-four hours post-transfection, cells were pre-treated with a Jak1/Jak2 inhibitor for one hour. Post pre-treatment, cells were mock- or SeV-infected in the presence of the inhibitor. Firefly and *Renilla* luciferase activities were determined eighteen hours later using a dual luciferase assay (Promega). Fold induction was determined relative to the DMSO vector only, mock-infected samples. Assays were performed in triplicate; error bars represent the SEM for the triplicate. Whole cell lysates (WCL) were analyzed by western blot with anti-FLAG and anti-β-tubulin antibodies. Statistical significance was determined by performing a one-way ANOVA followed with Tukey multiple comparison as compared to SeV-infected control (white bar); **p < 0.0001, * p < 0.001. **(B)** HEK293T cells were transfected with an ISG54 promotor-firefly luciferase reporter plasmid, a constitutively-expressing *Renilla* luciferase reporter plasmid, and empty vector. Twenty-four hours post-transfection, cells were pre-treated with DMSO or a Jak1/Jak2 inhibitor for one hour. Post pre-treatment, cells were mock- or UIFN-treated in the presence of the inhibitor. Firefly and *Renilla* luciferase activities were determined eighteen hours later using a dual luciferase assay (Promega). Fold induction was determined relative to the DMSO, mock-treated samples.

### MLAV VP24 fails to interact with Keap1 or activate ARE gene expression due to the absence of a Keap1-interacting K loop

MARV VP24 interacts with Keap1 to activate ARE promoters (36, 37). To determine whether MLAV VP24 possesses similar properties, co-immunoprecipitation experiments were performed with HA-tagged human Keap1 (hKeap1) or HA-tagged Keap1 derived from the bat *Myotis lucifugus* (bKeap1). As described, MARV VP24 interacted with both human and bat Keap1, whereas EBOV and MLAV VP24 did not (Figure 7A and 7B). Consistent with these data, when tested in an ARE promoter reporter gene assay, MARV VP24 activated the ARE reporter, relative to an empty vector control, while neither EBOV nor MLAV VP24 activated the ARE response. (Figure 7C).

**Figure 7.**
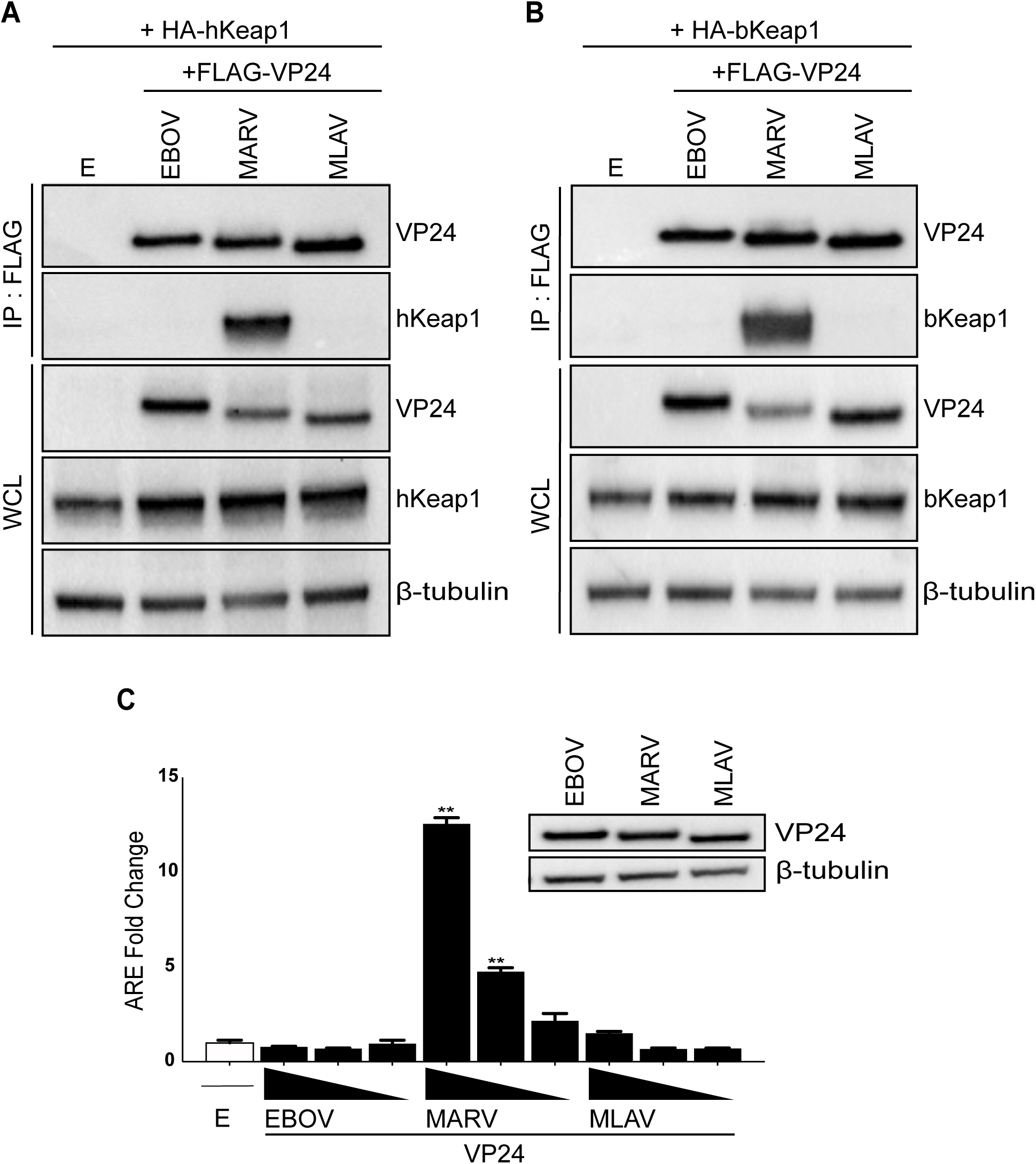
MLAV VP24 does not interact with KPNA5 or KEAP1. **(A-B)** HEK293T cells were co-transfected with FLAG-tagged EBOV, MARV, MLAV VP24, as indicated and HA-tagged human Keap1 (hKeap1) **(A)** or HA-tagged bat Keap1 (bKeap1) **(B)**. Co-immunoprecipitation (IP) was performed with anti-FLAG antibody and precipitates and whole cell lysates (WCL) were assessed by using anti-FLAG (VP24), anti-HA (Keap1) and anti-β-tubulin antibodies. **(C)** HEK293T cells were transfected with a reporter plasmid with the firefly luciferase gene under the control of an ARE promoter, a reporter plasmid that constitutively expresses *Renilla* luciferase and either empty vector (E) or the indicated FLAG-VP24 proteins. The concentration of VP24 plasmids were 62.5 ng, 6.25 ng and 0.625 ng. Firefly and *Renilla* luciferase activities were determined eighteen hours post-transfection. Firefly luciferase activity was normalized to *Renilla* luciferase activities and fold activity is reported, relative to the empty vector only sample. Protein expression was analyzed by western blot using anti-FLAG (VP24) and anti-β-tubulin antibodies. The assays were performed in triplicate, error bars represent the SEM for the triplicate. Statistical significance was determined by performing a one-way ANOVA followed with Tukey multiple comparison as compared to vector only control (white bar); **p < 0.0001, * p < 0.001.

MARV VP24 interaction with Keap1 occurs via a specific motif, the K-loop, and transfer of this sequence to EBOV VP24 confers binding to Keap1 (36). To determine whether this sequence could confer interaction with Keap1 and activation of ARE responses upon MLAV VP24, the MARV VP24 K-loop sequence (202-219) was transferred to MLAV VP24, replacing the corresponding amino acid residues (MLAV VP24_MARV 202-219_). The reverse chimera was also generated, with MLAV sequences replacing the K-loop in MARV VP24 (MARV VP24_MLAV 202-219_) (Figure 8A). Transferring the MARV K-loop sequence to MLAV VP24 conferred the capacity to activate an ARE response while transfer of the MLAV sequence to MARV VP24 abolished the activation (Figure 8B). Interaction with human Keap1 (Figure 8C) and bat Keap1 (Figure 8D) yielded corresponding data where interaction was dependent on the MARV VP24 K loop. Collectively, these data demonstrate that the lack of ARE gene expression by MLAV VP24 is due to the lack of a Keap1 binding motif.

**Figure 8.**
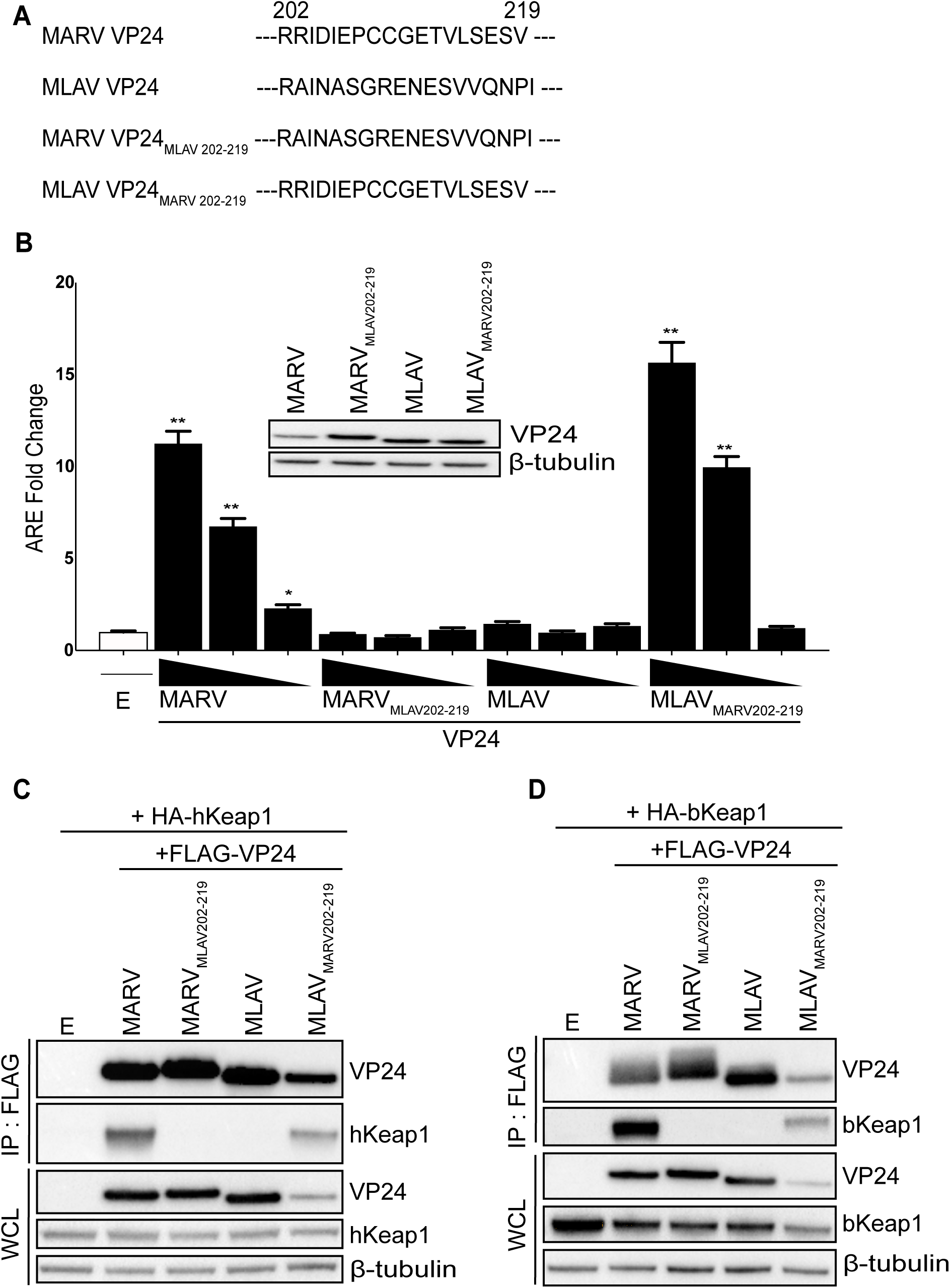
Transfer of the MARV K-Loop sequence confers on MLAV VP24 interaction with Keap1 and activation of ARE signaling. **(A)** Diagram of MARV and MLAV sequences for residues 202-219 and the VP24 chimera constructs MLAV VP24 _MARV 202-219_ and MARV VP24_MLAV 202-219_. **(B-C)** HEK293T cells were transfected with FLAG-tagged constructs, as indicated and either **(B)** HA-tagged human Keap1 (hKeap1) or **(C)** HA-tagged bat Keap1 (bKeap1). Co-immunoprecipitation (IP) was performed with anti-FLAG antibody. IPs were analyzed by western blotting with anti-FLAG (VP24), anti-HA (Keap1) and anti-β-tubulin antibodies. **(D)** HEK293T cells were transfected with reporter plasmid with the firefly luciferase gene under the control of an ARE promoter, a reporter plasmid that constitutively expresses *Renilla* luciferase and either empty vector (E) or the indicated FLAG-VP24 proteins. The concentration of VP24 plasmids were 62.5 ng, 6.25 ng and 0.625 ng. Firefly and *Renilla* luciferase activities were determined eighteen hours post transfection. Firefly luciferase activity was normalized to *Renilla* luciferase activities and fold activity is reported, relative to the empty vector only sample. Whole cell lysates (WCL) were harvested and lysates were analyzed by western blot with anti-FLAG (VP24) and anti-β-tubulin antibodies. The experiment was performed in triplicate, error bars represent the SEM for the triplicate. Statistical significance was determined by performing a one-way ANOVA followed with Tukey multiple comparison as compared to vector only control (white bar); **p < 0.0001, * p < 0.001.

## Discussion

The data in this study provide functional evidence that MLAV is biologically distinct from other filoviruses and support its classification in its own genus. The placement of MLAV in a distinct genus was based on its relatively low sequence identity to other filoviruses(1). It was also noted to have, compared to other filoviruses, unique gene overlaps and a unique transcription start signal. Despite these distinctions, MLAV mechanisms of entry and RNA synthesis mirror those of both EBOV and MARV. MLAV also possesses some features that suggest a closer genetic relationship to members of the *Marburgvirus* genus as opposed to the *Ebolavirus* and *Cuevavirus* genera. This includes similarities in Large (L) protein sequence and the absence of RNA editing sites in GP. In addition, MLAV was identified in *Rousettus* bats, and *Rousettus* bats in Africa serve as a reservoir for MARV and RAVV. The present study demonstrates the capacity of MLAV VP35 and VP40 to counteract IFN responses in human and bat cells. Inhibition of RIG-I induced IFN responses is thus far a common feature of filoviruses (52). The suppression of IFN-induced signaling and gene expression by VP40, rather than via VP24, parallels MARV and draws a functional distinction between MLAV and EBOV. The absence of MLAV VP24 interaction with human or bat Keap1, and its lack of ARE transcriptional activation is consistent with MLAV having evolved unique virus-host interactions that are distinct from MARV.

The data demonstrate that MLAV encodes mechanisms to counteract both IFN-α/β production and responses. MLAV VP35 was demonstrated to effectively block activation of the IFNβ promoter in response to SeV infection, a known inducer of the RIG-I signaling pathway. In addition, inhibition of SeV-induced phosphorylation of IRF3 was demonstrated. Together, these data indicate that MLAV can block RIG-I signaling, consistent with the function of other filovirus VP35s (53, 54). Mechanistically, inhibition of IFN-α/β production by EBOV or MARV VP35 correlates with dsRNA binding activity (9, 10, 12–14, 16, 19, 20, 45). This may reflect binding and sequestration of RIG-I activating dsRNAs (13, 20). The VP35 dsRNA binding domain, also known as the interferon inhibitory domain (IID), directly contacts the phosphodiester backbone of dsRNA, via residues that comprise a central basic patch, to mediate this interaction (10–12, 17, 18, 45). EBOV VP35 also caps the ends of dsRNA in a manner that likely masks 5’-triphosphates, which contribute to recognition of RNAs by RIG-I (12, 17). VP35 interaction with host protein PACT, which interacts with and facilitates activation of RIG-I, also contributes to inhibition (13, 55). Because the residues that make up the central basic patch are conserved between MLAV and other filoviral VP35s (1), MLAV is likely to bind to dsRNA. Given that it also interacts with PACT (Figure 1D), its mechanisms of inhibition are likely very similar to other filoviral VP35s.

EBOV, MARV and LLOV VP35 have also been demonstrated to inhibit activation of PKR, an IFN-induced, dsRNA-activated protein kinase that exerts antiviral effects by suppressing translation (21–24, 43). The mechanism by which VP35s inhibit PKR remains ambiguous, however, mutation of multiple central basic patch residues in EBOV or MARV VP35 disrupts the inhibitory activity (22, 23). In contrast, single point mutations that disrupt EBOV VP35 dsRNA binding activity leave PKR inhibition intact, suggesting that inhibition of PKR is not dependent upon VP35-dsRNA interaction or sequestration (21, 22). Consistent with PKR inhibition being an important function for filoviruses, this activity is conserved in MLAV as well.

The IFN-inhibitory activities of both EBOV and MARV VP35 have been demonstrated to be important for efficient virus replication in IFN-competent systems (14, 44). In addition to blocking the production of antiviral IFNs, VP35 inhibition of RIG-I also suppresses maturation of dendritic cells when expressed alone or in the context of EBOV infection (15, 56, 57). This activity impairs adaptive immunity to EBOV (58, 59). Therefore, VP35 likely inhibits adaptive, as well as innate, antiviral defenses. Disruption of VP35 anti-IFN function in the context of recombinant EBOVs has been demonstrated to render the virus avirulent in rodent models (14, 60). Based on these data, VP35 suppression of RIG-I signaling appears to be critical for virulence. The effective function in human cells of MLAV VP35 satisfies one apparent criteria for virulence in humans. It should be noted however, that suppression of RIG-I signaling by VP35 is not sufficient on its own to confer virulence. Even though MARV VP35 functions in *Rousettus* cells and likely has evolved in this species, MARV does not appear to cause significant disease in these animals (61–63). It does seem likely however, that in the reservoir host, VP35 IFN-antagonist function will be important for efficient replication and transmission, although this remains to be tested experimentally.

For MARV, either infection or VP40 expression alone blocks IFN induced phosphorylation of Jak kinases, inhibiting activation and downstream signaling. The absence of these phosphorylation events in response to IFN-α/β or IFNγ is consistent with the phenotype of Jak1-deficient cells, suggesting that Jak1 function may be targeted by MARV VP40, although there is no evidence to date of VP40-Jak1 interaction (31). Consistent with MARV VP40 impairing Jak1 function, MARV VP40 expression is sufficient to prevent phosphorylation of STAT proteins following Jak1 over-expression or treatment by IFN-α/β or IFNγ (type II IFN) (31). MLAV VP40 likewise blocks ISG expression and inhibits STAT1 phosphorylation following IFN treatment or over-expression of Jak1. Therefore, inhibition of IFN signaling by MLAV VP40 seems likely to proceed by a mechanism similar to that employed by MARV VP40.

MARV VP24 binds directly to Keap1, a cellular substrate adaptor protein of the Cullin-3/Rbx1 E3 ubiquitin ligase complex (36–38, 64). Keap1 regulates the cellular antioxidant response (65). Under homeostatic conditions, Keap1 promotes Nrf2 polyubiquitination and degradation However, cell stresses, including oxidative stress, disrupt the Keap1-mediated ubiquitination of Nrf2, stabilizing it and promoting Nrf2 dependent expression of antioxidant response genes. Biophysical studies demonstrated that MARV VP24 interacts with the Keap1 Kelch domain at a site that overlaps the region that binds Nrf2 (38). This interaction disrupts Nrf2-Keap1 interaction and activates ARE gene expression (36–38). Keap1 similarly interacts with host kinase IKKβ to repress NF-ĸB responses and MARV VP24 can also disrupt this interaction, thereby relieving Keap1 repression on the NF-ĸB transcriptional response (39). In contrast, EBOV and LLOV VP24 targets KPNA proteins in a manner that prevents pY-STAT1 nuclear transport, inhibiting ISG expression (25, 26, 29, 30, 43).

Given that MLAV VP40 mirrored MARV VP40 in its inhibition of the IFN response, it was of interest to determine whether MLAV VP24 would similarly mimic MARV VP24 in terms of interaction with host signaling pathways. However, MLAV VP24 lacks a sequence that resembles the MARV VP24 K-loop and, correspondingly, did not interact with human or a bat-derived Keap1 and did not activate an ARE promoter. Chimeric MARV-MLAV VP24 proteins confirmed that the absence of the K-loop sequence can explain the lack of MLAV VP24 effects on antioxidant responses. Furthermore, consistent with the absence of MLAV VP24 inhibitory activity in IFN-signaling assays, it also fails to interact with KPNA5. The interface between EBOV VP24 and KPNA covers a large surface area and involves multiple points of contact (30). This precluded the mapping of specific amino acid residues that explain the lack of MLAV VP24-KPNA5 interaction. Nonetheless, these data presented here indicate that MLAV VP24 does not reflect the functions of either MARV or EBOV VP24. It will be of interest to determine whether MLAV VP24 engages different host pathway(s).

The inhibition of IFNβ promoter activity by MLAV VP40 parallels the inhibition by EBOV VP24 and MARV VP40, although inhibition by MLAV VP40 appeared to be more potent. MARV VP40 and EBOV VP24 inhibition of IFN-α/β production and, in the case of EBOV VP24, production of IFN-λ as well, have been previously reported (34, 35). However, the mechanism(s) for these inhibitory activities are incompletely defined, although EBOV VP24 was implicated as having an effect post-IRF3 phosphorylation (34). Inhibition of STAT1 activation and IFN-induced gene expression would be expected to impair the positive feedback loop in which IFN-α/β induces expression of IFN stimulated genes, including RIG-I and IRF7, to amplify IFN response (51). This prompted additional experiments to determine whether the detected inhibition was a product of blocking the positive feedback loop through VP40 inhibition of Jak-STAT signaling or an additional mechanism acting on the production side. Upon treatment of cells with a Jak1/Jak2 inhibitor, which removes any potential contribution of IFN signaling inhibition, MLAV VP40 maintained inhibition of the IFNβ promoter. This suggests MLAV VP40 has an additional mechanism(s) of IFN antagonism that requires further exploration.

Cumulatively, the present study has identified several functions of MLAV proteins that, in conjunction with previously published data, indicate a compatibility with infection of humans. These include the capacity of MLAV GP to mediate entry into human cells via interaction with NPC1 and suppression of IFN responses through several mechanisms (1). Notably, given that MLAV VP24 does not detectably interact with either KPNA5 or Keap1, it is likely that it may make unique interactions with host cells, having unknown contributions to the regulation of cellular signaling pathways. Therefore, the existing data also suggests that the outcome of MLAV infection in humans could differ from that of the typical outcome of EBOV or MARV infection.

## Acknowledgements

The following reagent was obtained through BEI Resources, NIAID, NIH: RO6E, *Rousettus aegyptiacus* (Egyptian fruit bat), Immortalized Fetal Cell Line, NR-49168. This work was supported by the following grants to C.F.B.: NIH grant AI120943, NIH grant AI109945 and by Department of the Defense, Defense Threat Reduction Agency grant HDTRA1-16-1-0033 (C.F.B.). The content of the information does not necessarily reflect the position or the policy of the federal government, and no official endorsement should be inferred. C.F.B. is a Georgia Research Alliance Eminent Scholar in Microbial Pathogenesis.

